# MicroRNA-9 Mediated the Protective Effect of Ferulic Acid on Hypoxic-Ischemic Brain Injury in Neonatal Rats

**DOI:** 10.1101/2020.01.27.920900

**Authors:** Keli Yao, Qin Yang, Yajuan Li, Ting Lan, Hong Yu, Yang Yu

## Abstract

Neonatal hypoxic-ischemic brain damage (NHIBD) leads to cognitive and memory impairments, and there is no effective clinical treatment. Ferulic acid (FA) is found within members of the genus *Angelica*, reportedly shows protective effects on neuronal damage; however, the mechanism of the protective effects of FA on rats following NHIBD remains unclear. Using the Morris water maze task, we herein found that the impairment of spatial memory formation in adult rats exposed to NHIBD was significantly reversed by FA treatment and the administration of LNA-miR-9. RT-PCR analyses revealed that miRNA-9 was significantly increased in the hippocampus of neonatal rats and neuronal PC12 cells following NHIBD and that FA and LNA-miR-9 both inhibited the expression of miRNA-9, suggesting that the therapeutic effect of FA was mainly attributed to the inhibition of miRNA-9 expression. Indeed, the silencing of miR-9 by LNA-miR-9 or FA similarly attenuated neuronal damage and cerebral atrophy in the rat hippocampus after NHIBD, which was consistent with the restored expression levels of brain-derived neurotrophic factor (BDNF). Therefore, FA treatment may protect against neuronal death through the inhibition of miRNA-9 induction in the rat hippocampus following hypoxic-ischemic damage.

## Introduction

Neonatal hypoxic-ischemic brain damage (NHIBD) is a severe disease that can cause irreversible neurological sequelae, such as cerebral palsy, mental deficiency, memory impairment and learning disabilities and is often characterized by permanent neurological deficits [1,2]. Currently, there is no effective clinical treatment for NHIBD [3–5]. Thus, effective therapeutic agents that inhibit damage cascades activated after NHIBD should be identified. MicroRNAs have recently been identified as critical mediators of neuroinflammation and neurodegeneration in NHIBD [6,7]. Recent reports have demonstrated that miR-30b is upregulated during NHIBD and is involved in the regulation of cellular apoptosis after injury [8]. MicroRNA-9 (miR-9) is highly expressed in the brain and has been implicated in the regulation of neurogenesis and proliferation as well as axonal development and neuronal migration [9–13]. However, little attention has been paid to the role of miR-9 in brain damage in infants caused by hypoxia.

Ferulic acid (FA) is a widely distributed constituent of plants that can treat cerebrovascular diseases and cerebral ischemia by protecting neurons from degeneration and regulating inflammatory activity [14–16]. However, it is unknown whether the administration of FA has a neuroprotective effect on HIBD; furthermore, the potential mechanisms remain unclear. Therefore, the purpose of this study was to further characterize the neuroprotective effect of FA on learning and memory ability, possibly via the downregulation of miR-9 following hypoxic-ischemic injury.

## 2. Methods and materials

### 2.1 Preparation of SF and LNA-miR-9

Sodium ferulic (SF, the sodium salt form of FA, purity >99.9%) was purchased from Dalian Meilunbio Biotechnology Co., Ltd. (CAS Registry Number: 24276-84-4, China), dissolved in 0.9% saline solution to a final concentration of 7 mg/ml, and subsequently stored at 4°C.

The sequence of LNA-miR-9 is 5’-tcatacaGCtAgAtaACcaAaGa-3’, with the uppercase letters representing the sites of locked nucleic acid modifications [17]. LNA-miR-9 was synthesized by Sangon Biotech Co., Ltd. (Shanghai, China), dissolved in PBS to a final concentration of 200 µM and stored at −20°C.

### 2.2 Cell culture and HI model establishment

PC12 cells were incubated in a humidified atmosphere (95% air and 5% CO_2_) at 37°C. Fresh medium was supplied every 48 h. Cultured PC12 cells were serum-starved in RPMI-1640 medium supplemented with 1% fetal calf serum and 1% horse serum and placed at 37°C in a humidified three-gas incubator (2% O_2_, 93% N_2_, and 5% CO_2_; NUAER) for 2 h to induce hypoxic-ischemic injury [18]. Then, the cells were treated with sodium ferulic (SF) or LNA-miR-9 for 24 h.

### 2.3 Animals and HIBD model establishment

Seven-day-old Sprague-Dawley (SD) rats were provided by the Animal Department of Southwest Medical University (license number: SYXK (Sichuan) 2018-065, LuZhou City, China). An HIBD model was established by the classic method of Rice-Vannucci. The left carotid artery of 7-day-old SD rats was permanently unilateral ligated and removed, and the pups were returned to their mothers for 1 h for recovery. Then, the pups were exposed in a hypoxic environment (8% O_2_ and 92% N_2_) for 2 h [19,20]. In sham animals, the vagus nerve was separated but not ligated, and the pups were placed in a similar container but exposed to normal room air. All experiments were performed in compliance with the Health Guide for the Care and Use of Laboratory Animals and approved by Sichuan Experimental Animal Management Committee.

### 2.4 Drug treatment

Immediately after the HIBD model was established, when the pups were 21 days old, 50 mg/kg SF was intraperitoneally injected into the rats in the SF group one time per day for 5 days. For the LNA-miR-9 group, the rat received a terminal intraperitoneal injection of sodium pentobarbital, 2 μl (400 μM) of LNA-miR-9 was stereotaxically microinjected into the lateral ventricle within 15 min. The dwell time was 5 min, and the injector was positioned 1.0 mm posterior to bregma, 0.8 mm from the midline, and 3.5 mm deep from the dura. The skin wound was sutured after the surgery. LNA-miR-9 treatment was only administered for one day. Considering that the robust neuroprotective effects of SF (0.5 µg/ml) and LNA-miR-9 (50μM/ml) were observed, we decided to use those concentration in the following *in vitro* studies.

### 2.5 Real-time PCR

Total RNA was extracted from the rat hippocampus after various treatments using an RNA-simple Total RNA Kit (TIANGEN, China) and then reverse transcribed using a ReverTra Ace qPCR RT Master Mix Kit (TOYOBO, Japan) and a miRcute Enhanced miRNA cDNA First-Strand Synthesis Kit (TIANGEN) according to the manufacturers’ protocols. Real-time PCR was performed on a qTOWER^3^G (Analytikjena, Germany). The specific PCR primers use for the detection of acetylcholinesterase (AchE), postsynaptic density protein-95 (PSD-95) and β-actin were designed according to the NCBI sequence and synthesized by Sangon Biotech Co., Ltd. The sequences of the primers were as follows:

**Table 1.**
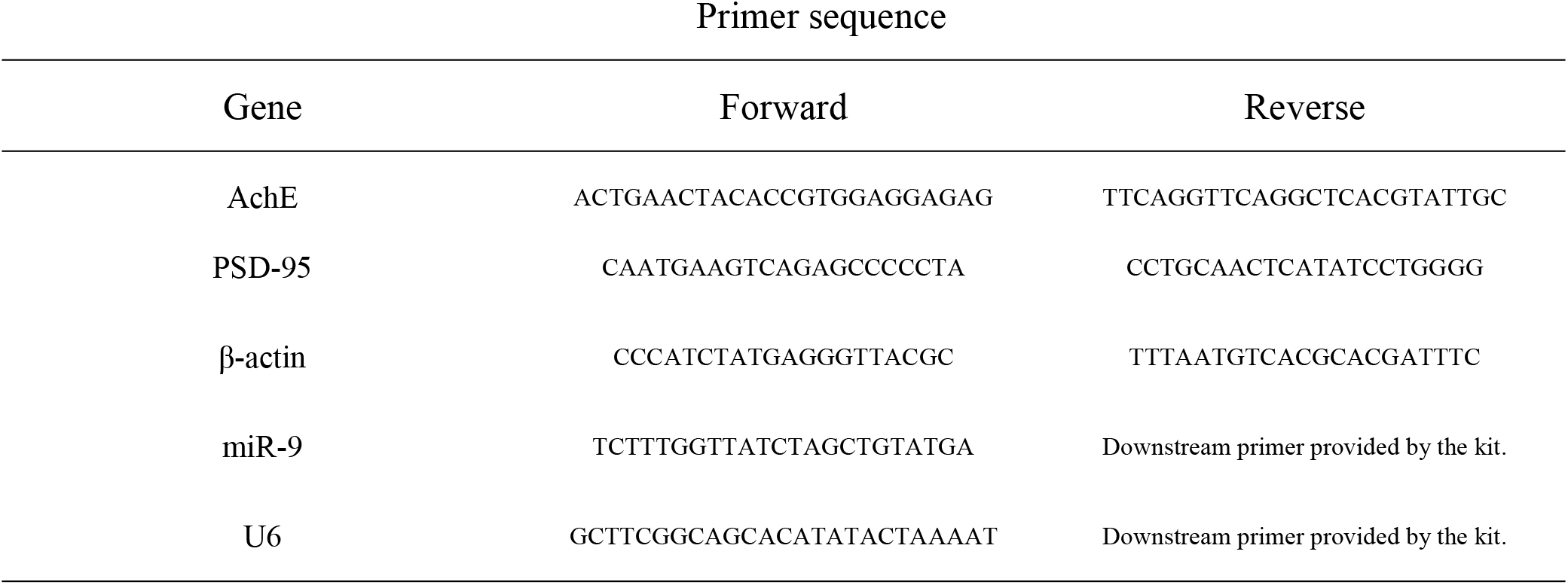
Primer sequence

### 2.6 Western blotting

Western blotting was performed as described previously [21]. In brief, PC12 cells and hippocampal tissues were washed with ice-cold PBS three times and then lysed in lysis buffer containing protease inhibitor tablets. Total protein was extracted for Western blotting. Then, total proteins were separated on 10% SDS-PAGE gels and transferred onto PVDF membranes. The membranes were blocked with 5% skim milk solution in TBS-T for 1 h at room temperature. After blocking, the membranes were incubated with primary antibodies (anti-AchE, 1:1000, Abcam, USA; anti-PSD95, 1:3000, Abcam, USA; anti-β-actin, 1:10000, Proteintech, China) overnight at 4°C. The membranes were washed three times with TBS-T and then incubated with secondary antibodies (1:3000, Proteintech, China) for 1 h at room temperature. After washing, the membranes were placed into chemiluminescent HRP substrate (Millipore, USA) for 2 mins, and the protein bands were detected with a Clinxchemi Scope 6000 (Clinx, China). The absorbance values of the bands were quantified using an image analysis system and Image J. The protein levels were normalized against the β-actin intensity.

### 2.7 H-E staining

Hematoxylin-eosin (H-E) staining was performed on paraffin sections of brain tissue. Anesthetized rats were perfused with saline followed by 4% paraformaldehyde solution. The brains were then collected and immersed in 4% paraformaldehyde. After dehydration and vitrification, the brains were embedded in paraffin and cut into 5-6-μm sections. The sections were adhered to glass slides precoated with polyethylenimine. Paraffin-embedded brain sections were dewaxed with xylene and rehydrated with an ethanol series (absolute, 95%, 90%, 80%). The sections were then stained with H-E, and images were collected under a microscope.

### 2.8. Morris water maze test

Four weeks after HIBD, 30-day-old rats were subjected to the Morris water maze test as described previously [22]. In brief, an open circular water-filled pool with a diameter of 160 cm, a height of 50 cm and a temperature of 22±1°C was prepared. The pool was divided into four quadrants, each of which had a reference object consisting of a different colored shape. In the experiment, an escape platform below the water surface was placed in a quadrant, rats were placed into the pool and allowed to swim, and the amount of time needed and the route used to find the platform were recorded. The swimming behavior of the rats was recorded and tracked by an overhead video camera connected to a PC with an automated tracking software, and the data were analyzed using the behavior analysis system (Shanghai Xinruan Information Technology Co., Ltd., Shanghai, China).

### 2.9 Statistical Analyses

All data are expressed as the mean ± standard deviation. Statistical analyses were performed using GraphPad Prism 7 software. Differences between two groups were evaluated statistically by using unpaired Student’s t test. One-way ANOVA was used to determine the significance of the differences among the experimental groups. *P*< 0.05 was regarded as the level of statistical significance.

## 3. Result

### 3.1 FA ameliorated cerebral palsy, memory impairment and learning disabilities in HIBD rats

To determine whether FA can rescue HIBD-induced cerebral palsy, memory impairment and learning disabilities in neonatal rats, H-E staining and the Morris water maze test were conducted. H-E staining showed that the number of hippocampal neurons in the CA1 area, the size of the hippocampus and the layers of the dentate gyrus were dramatically decreased in the HIBD group compared with the sham group and that the administration of FA significantly suppressed the decrease in hippocampal neurons and the increases in the size of the hippocampus and the layers of the dentate gyrus (Fig. 1A-C, sham: N = 6, HIBD: N = 6, HIBD + SF: N =6, *P* < 0.001 vs. HIBD).

**Fig 1.**
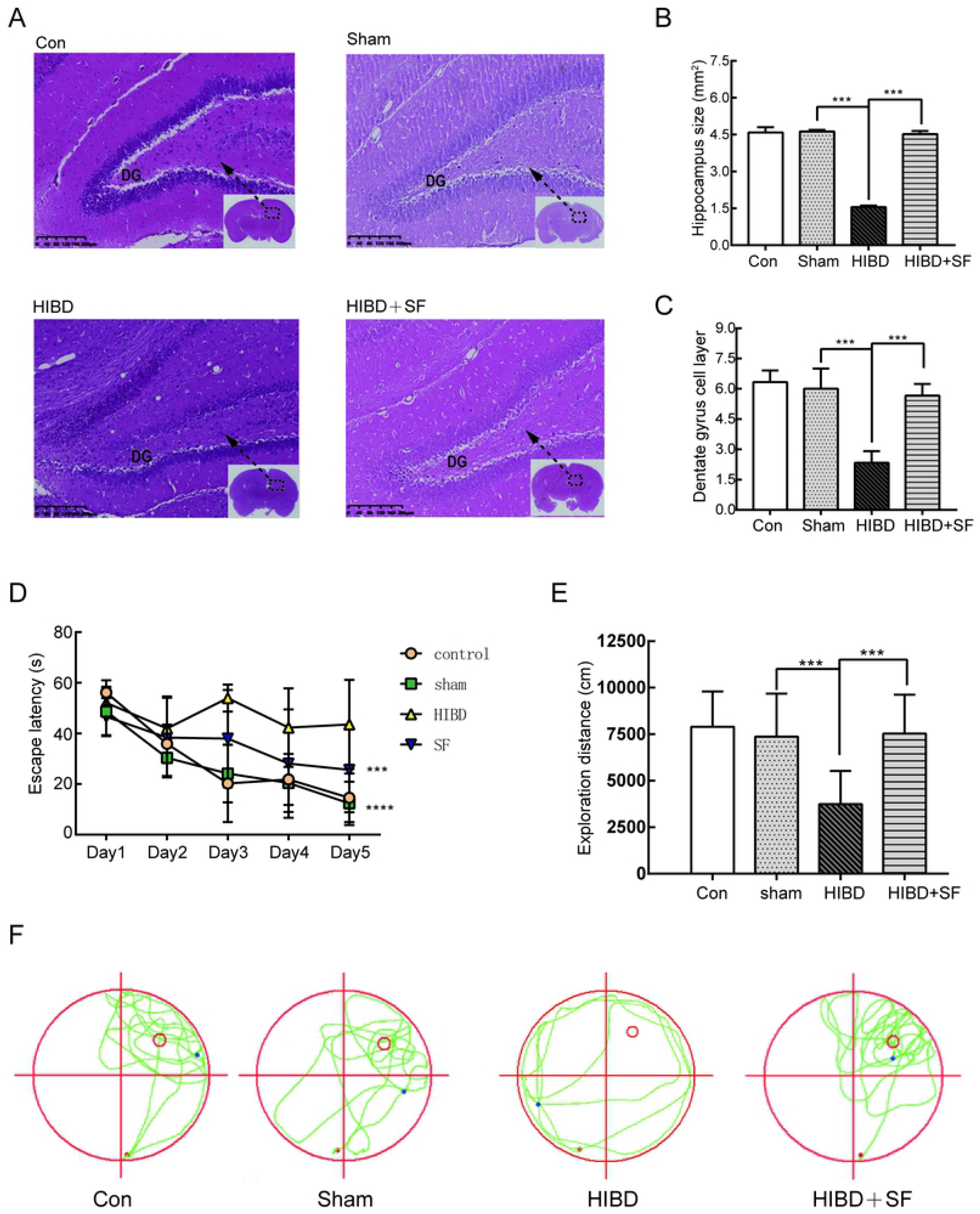
FA ameliorated brain tissue loss and learning and memory impairments. (A) Coronal sections stained with hematoxylin/eosin are shown. (B, C) Quantification of the size of the hippocampus and the layers of the hippocampal dentate gyrus in the different groups. (D) The escape latency (T value) of the rats in the training period. (E) The exploration distance (P value) of the rats in the exploratory trials. (F) The exploration route in the retention trial. *\*\*\* P* < 0.001*, ** P* < 0.01. N=6 for H-E staining and N=10 for Morris water maze analysis.

To evaluate long-term spatial learning and memory ability, the Morris water maze was performed on 35-day-old rats after SF application. In the acquisition trial, HIBD rats showed decreased spatial learning ability with a significantly increased escape latency compared with those of the control and sham groups on the training days. Importantly, as shown in Fig. 1D, daily SF treatment (50 mg/kg, i.p.) significantly ameliorated the impairment induced by HIBD (Fig 1D, *P* <0.001). In the retention trial, a significant reduction in the exploration distance in the target quadrant (Fig 1E-F, *P* < 0.001) was observed in HIBD rats compared with HIBD+SF rats (*P* < 0.001). These results further confirmed that spatial memory retrieval was impaired after HIBD and that FA treatment succeeded in preventing this impairment.

Taken together, these results indicate that FA treatment alleviates deficits in ischemia-induced cell death and learning and memory function in neonatal rats after HIBD.

### 3.2 The expression of miR-9 significantly increased under hypoxic-ischemic conditions, and FA attenuated the expression of miR-9

To establish whether miR-9 is involved in HIBD, we investigated the expression levels of miR-9 mRNA in the hippocampus using qRT-PCR. The qRT-PCR results showed that the expression level of miR-9 in PC12 cells was markedly higher after HI injury than in the control groups (Fig 2A, *P* <0.05). Similarly, miR-9 expression was almost doubled in the hippocampus of hypoxic-ischemic rats compared to sham rats (Fig 2B, *P* <0.01). To further determine the therapeutic effects of FA on HIBD, we examined the expression levels of miR-9 after treating the rats with SF daily for 5 days and then compared them with the levels in the sham group. In PC12 cells cultured under serum-oxygen deprivation conditions for 2 h, miR-9 release was significantly reduced after treatment with SF (Fig 2C, *P* <0.05). Similar results were also observed in HIBD rats, and the expression levels of miR-9 were markedly lower in the HIBD+SF group than in the HIBD group (Fig 2D, *P* <0.01). These results further confirmed that miR-9 plays an important role in neonatal HIBD and that the neuroprotective effect of FA may occur through the downregulation of the expression levels of miR-9.

**Fig. 2.**
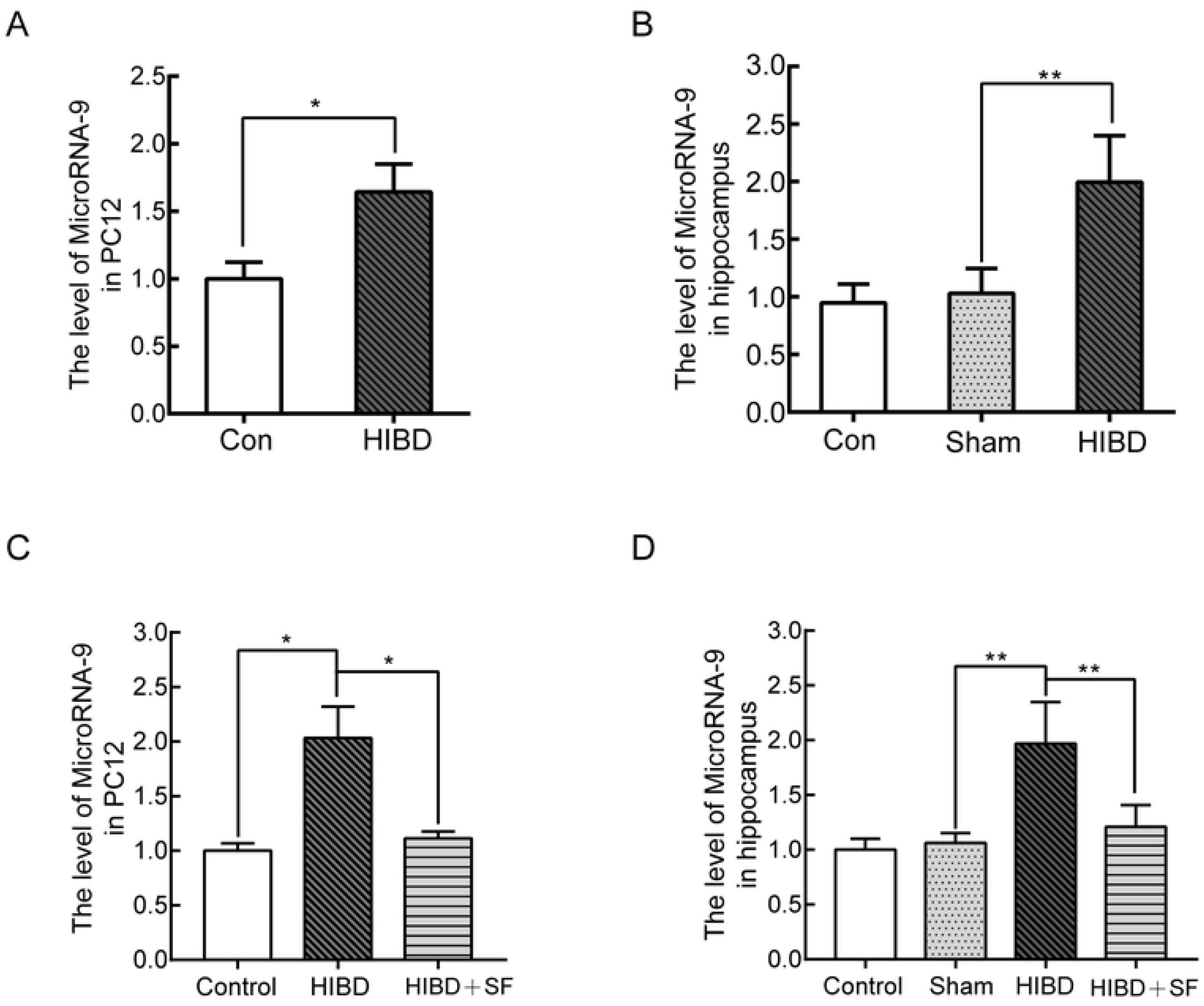
FA attenuated the upregulation of miR-9 under hypoxic-ischemic conditions in cultured PC12 cells and the rat hippocampus. (A) RT-PCR analysis of miR-9 expression in PC12 cells after HI. (B) RT-PCR analysis of miR-9 expression in the rat hippocampus after HIBD. *\*\* P* < 0.01*, * P* < 0.05 vs. control. (C) RT-PCR analysis of miR-9 expression in PC12 cells after FA treatment. (D) RT-PCR analysis of miR-9 expression in the rat hippocampus after FA treatment. *\*\* P* < 0.01*, * P* < 0.05 vs. HIBD. N= 4 for cell analysis and N= 5 for animal analysis.

### 3.3 Inhibition of miR-9 protected against hypoxic-ischemic brain injury in neonatal rats

To further reveal the role of miR-9 following hypoxic-ischemic damage. We investigated whether inhibiting the expression of miR-9 in vivo protects hippocampal neurons. We injected LNA-miR-9 (400 μM) into the lateral ventricle of 21-day-old neonatal rats after HIBD. LNA-miR-9 treatment significantly reduced brain injury in the rats, which resulted in a significant increase in the surviving brain volume and cell layers of the dentate gyrus compared to that of the HIBD+PBS group (Fig 3A-C,*P* <0.001). The Morris water maze was performed on 35-day-old rats after LNA-miR-9 application.

**Fig. 3.**
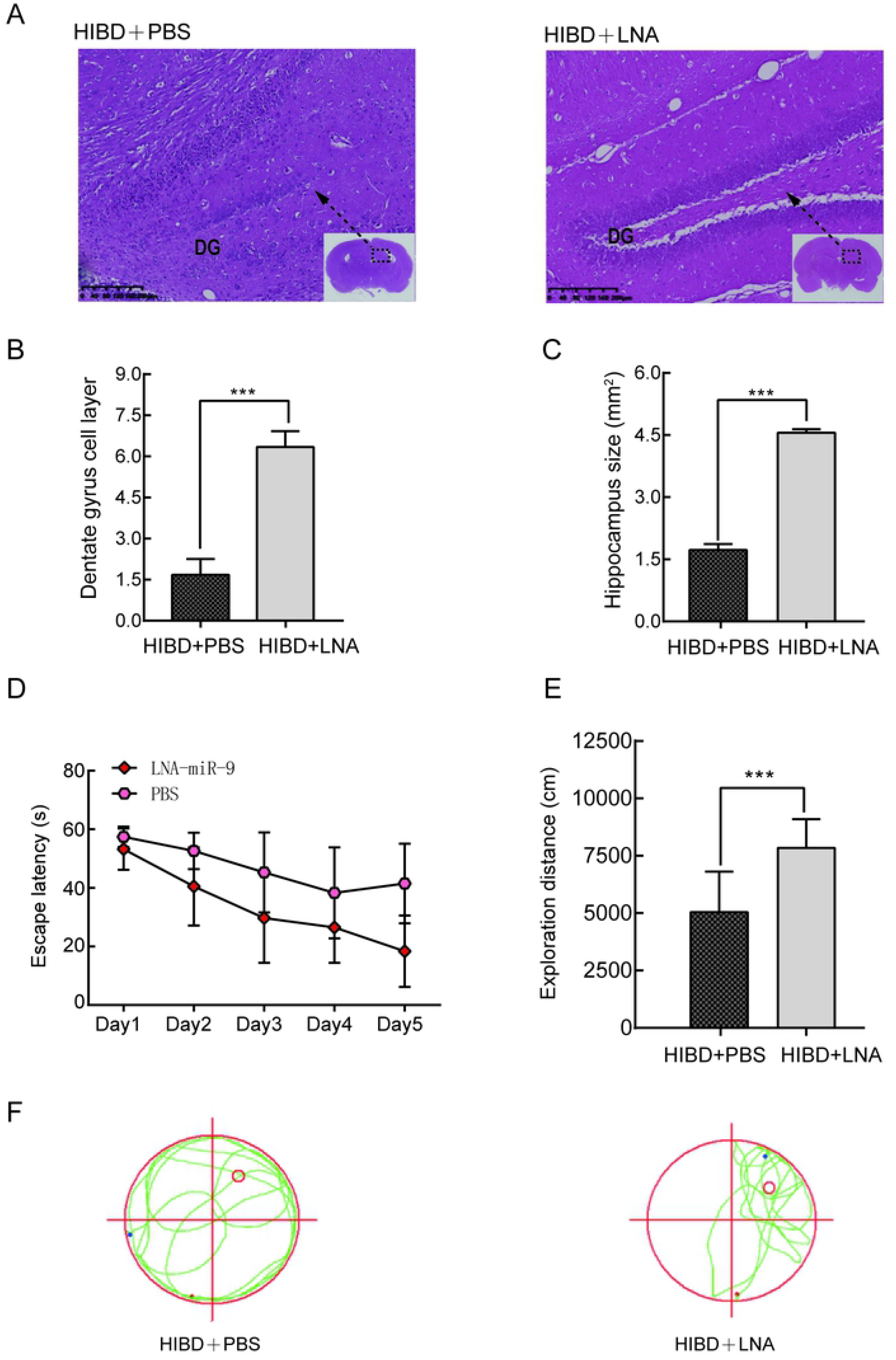
LNA-miR-9 attenuated brain tissue loss and learning and memory impairments. (A) Coronal sections stained with hematoxylin/eosin are shown. (B, C) Quantification of the size of the hippocampus and the layers of hippocampal dentate gyrus in the different groups. (D) The escape latency (T value) of the rats in the training period. (E) The exploration distance (P value) of the rats in the exploratory trials. (F) The exploration route in the retention trial. *\*\*\* P* < 0.001*, ** P* < 0.01 vs HIBD+PBS. N=6 for H-E staining and N=10 for Morris water maze analysis.

In the acquisition trial, HIBD+PBS rats showed decreased spatial learning ability with a significantly increased escape latency (Fig 3D, *P* < 0.001). Notably, LNA-miR-9 treatment significantly ameliorated the impairments of HIBD+PBS rats (Fig 3D, *P* <0.001). In the retention trial, a significant reduction in the exploration distance in the target quadrant (Fig 3E-F, *P* < 0.001) was observed in HIBD rats compared with HIBD+SF rats (*P* < 0.001). These results indicated that LNA-miR-9 protected against brain injury induced by HIBD in rats.

### 3.4 FA and LNA-miR-9 protected against synaptic marker loss induced by HI

Acetylcholinesterase (AchE) is a polymorphic enzyme commonly known for its cholinergic action that stops cholinergic signaling in the brain by hydrolyzing acetylcholine to choline and acetate [23]. The expression of AchE is closely correlated with the neuroinflammatory response and neurological dysfunction [24,25]. Studies have shown that the expression of AchE is increased after hypoxia-ischemia (HI) in newborn rats, and further research has confirmed that the inhibition of the AchE receptor reduces brain damage after HI [26]. In addition, previous reports have indicated that postsynaptic density protein (PSD)-95 expression is markedly decreased in the hippocampal CA1 region after brain injury; therefore, we measured the expression of PSD-95 and AchE to evaluate functional recovery following HIBD [27–29].

As shown in Fig 4A-B, the expression of AchE mRNA in the rat hippocampus and PC12 cells was significantly higher in the HIBD group than in the control group (*P <*0.01). SF and LNA-miR-9 notably decreased the HIBD-induced upregulation of AchE mRNA (*P <*0.05 vs HIBD). On the other hand, hypoxia-ischemia significantly reduced the expression of PSD-95 mRNA (*P <*0.05 vs control). After SF and LNA-miR-9 treatment, the expression of PSD-95 mRNA increased (Fig 4C-D, *P* <0.05 vs HIBD).

**Fig. 4.**
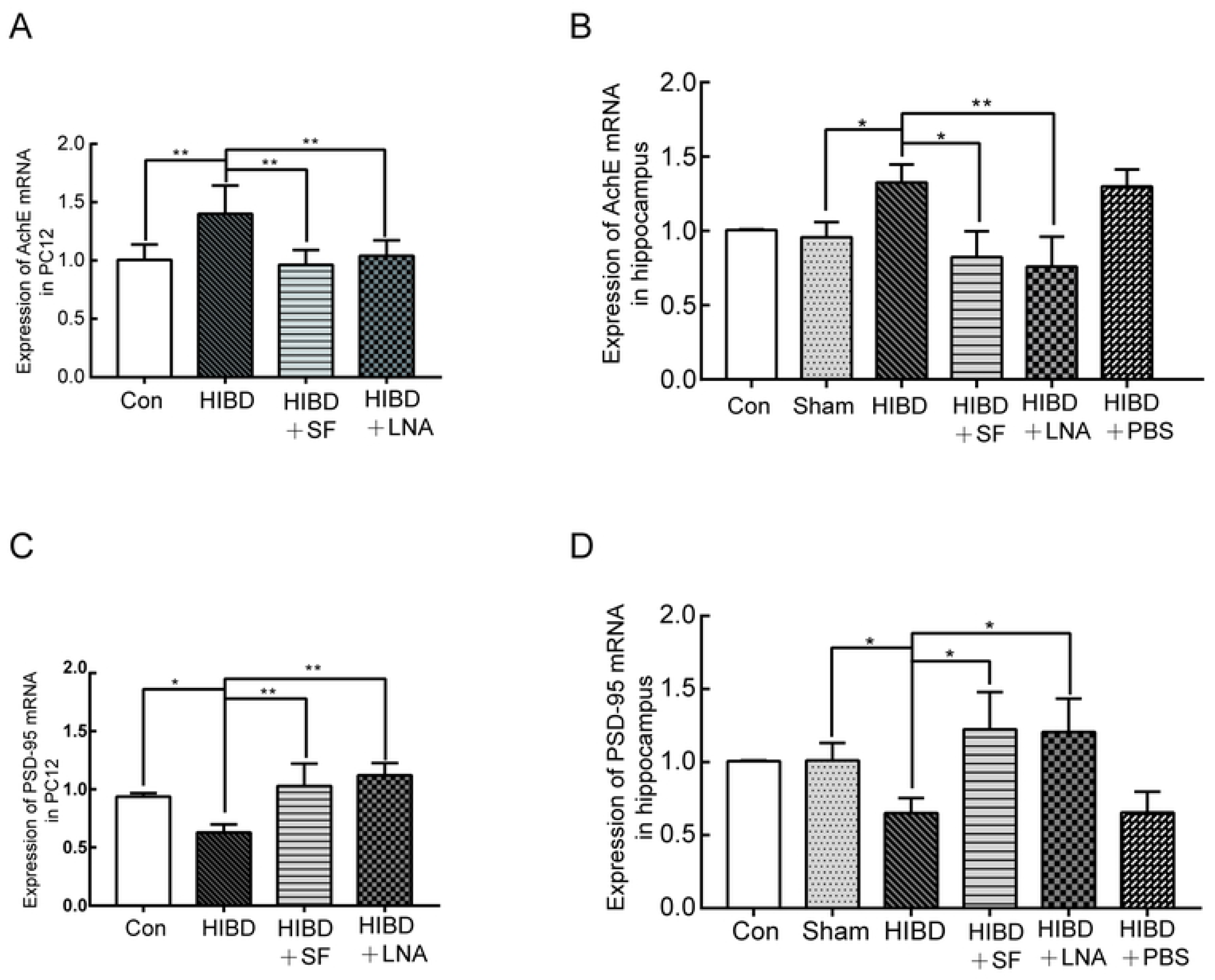
RT-PCR analysis of the gene expression of AchE and PSD-95 in PC12 cells and rat hippocampus. SF and LNA effectively reversed the expression changes in AchE and PSD-95 caused by hypoxic-ischemic injury. (A) The expression of AchE mRNA in PC12 cells. (B) The expression of AchE mRNA in the rat hippocampus. (C) The expression of PSD-95 mRNA in PC12 cells. (D) The expression of PSD-95 mRNA in the rat hippocampus. *\*\* P < 0.01, * P < 0.05*. N= 5 for cell analysis and N= 6 for animal analysis.

In addition, Western blot analysis was performed to verify the protein expression levels of AchE and PSD-95 in the rat hippocampus and PC12 cells. The results shown in Fig 5A-B and Fig 5C-D suggest that hypoxia-ischemia significantly increased the protein levels of AchE and decreased the levels of PSD-95 compared with those in the control group (*P* <*0.05*). Meanwhile, treatment with SF and LNA-miR-9 reduced the protein levels of AchE and upregulated the protein expression of PSD-95 (*P <0.05* vs HIBD). Collectively, these results indicated that miR-9 mediated the neuroprotective effect of FA against HIBD.

**Fig. 5.**
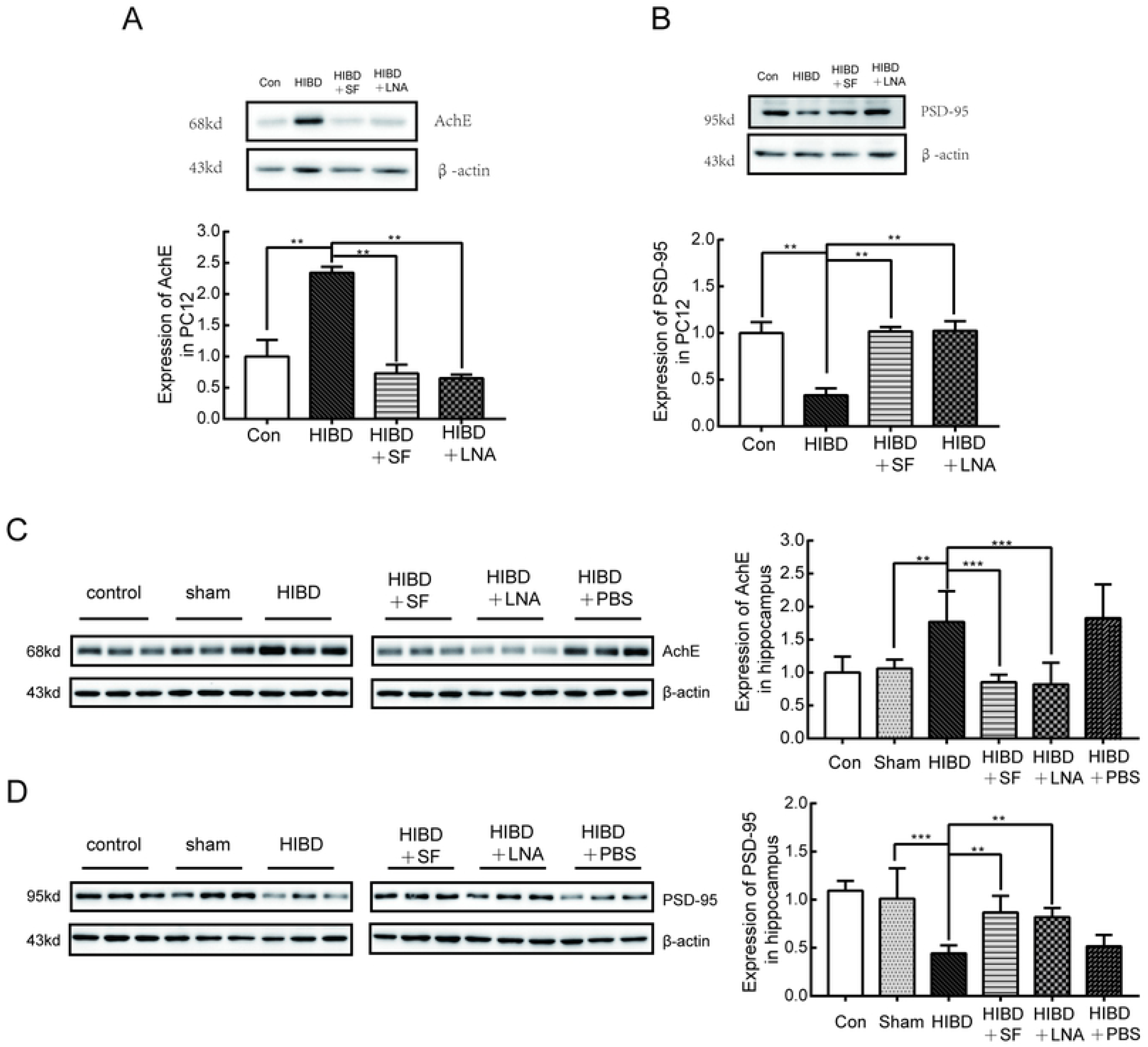
Western blot analysis of the protein expression of AchE and PSD-95 in PC12 cells and rat hippocampus. SF and LNA-miR-9 effectively reversed the expression changes in AchE and PSD-95 caused by ischemic-hypoxic injury. (A) The expression of AchE in PC12 cells. (B) The expression of AchE in the rat hippocampus. (C) The expression of PSD-95 in PC12 cells. (D) The expression of PSD-95 in the rat hippocampus. *\*\* P* < 0.01*, * P* < 0.05 vs HIBD or HIBD+PBS. N= 5 for cell analysis and N= 6 for animal analysis.

### 3.5 BDNF signaling participated in the neuroprotective effects of FA in HIBD neonatal rats

Brain-derived neurotrophic factor (BDNF) has been recognized as a protective effector of synaptic transmission and cognitive function after HI brain injury [30]. Previous reports have shown that the expression levels of BDNF are directly regulated by miRs and that miR-9 plays an important role in the regulation of axonal projections in response to BDNF signaling [31]. Therefore, we examined whether BDNF signaling functions in mediating miR-9 expression in HIBD neonatal rats. Western blot analysis revealed that BDNF expression was significantly decreased in the hippocampus of the rats in the HIBD group compared with those in the sham group after hypoxic-ischemic damage (*P* <0.01). In the neonatal rats in the HIBD+SF and HIBD+LNA-miR-9 groups, increased BDNF protein expression was observed (Fig 6A, *P* <0.01). These results confirmed that FA regulated miR-9 expression by increasing the release of BDNF.

**Fig. 6.**
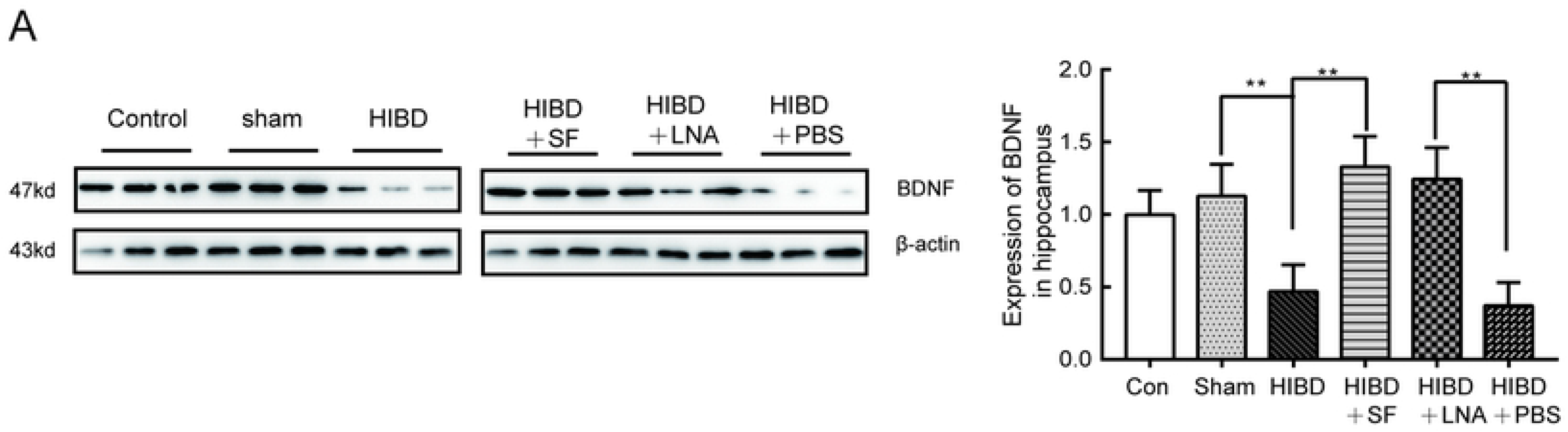
Western blot analysis of the protein expression of BDNF in the rat hippocampus. SF and LNA-miR-9 effectively increased the expression of BDNF in HIBD. (A) The expression of BDNF in rat hippocampus. ** *P* < 0.01 vs HIBD or HIBD+PBS. N= 6.

## Discussion

HIBD is one of the major causes of neonatal death worldwide [32]. It is urgent to identify drugs that protect against neuronal damage induced by HIBD. In this study, we demonstrated for the first time that FA can improve learning and memory impairment caused by HIBD in rats. The neuroprotective effects of FA are likely the result of the regulation of miR-9 expression through increased release of BDNF.

A growing number of researchers have discovered the beneficial influence of FA in stress-induced injury [33]. Treatment with FA has a protective effect by reducing oxidative stress and inhibiting ROS production in PC12 cells treated with the neurotoxin lead acetate [34]. Consistently, a previous report also revealed that FA possesses the ability to inhibit LPS-induced neuroinflammation [35]. However, little is known about its protective effects in HI brain injury. Previous reports have indicated that hypoxia-ischemia can reduce the long-term spatial learning and memory ability of rats [36]. Behavioral experiments showed that after treatment with SF and LNA-miR-9, the T value was significantly shortened and the P value was significantly prolonged compared with those of the HIBD group, suggesting that FA can improve the spatial learning and memory ability of HIBD rats.

MicroRNAs are small, noncoding RNAs that negatively regulate the expression of target mRNAs through degradation and translational inhibition [37]. An increasing number of studies have shown that microRNAs play an important regulatory role in neurodevelopment, metabolism and cancer [38–40]. miR-9 is a highly expressed miRNA in the brain and is enriched in synapses. Studies have shown that miR-9 can promote proliferation, migration, differentiation and apoptosis and regulate the dendrites of neurons [41–44]. However, whether miR-9 is involved in hypoxic-ischemic injury and recovery in the rat brain has not been clearly reported. This study found that the expression of miR-9 in PC12 cells and the hippocampus of the rats in the model group was significantly upregulated compared with that in the normal group, indicating that ischemia-hypoxia can influence the level of miR-9 expression in the process of neuronal damage. In addition, after SF treatment, the expression of miR-9 was significantly lower, revealing that FA may improve the learning and memory ability of HIBD rats by downregulating the expression of miR-9.

BDNF is broadly expressed in the developing mammalian brain, and evidence has shown that hypoxia-ischemia in rats results in increased BDNF expression in neurons [45]. Our experiments found that BDNF expression was significantly increased after treatment with FA and the inhibition of the expression levels of miR-9 in HIBD. In addition, the inhibition of miR-9 can downregulate the expression of AchE and upregulate the expression of PSD-95. The mechanism may be that the increased release of BDNF can protect against neuronal death in the rat hippocampus. This ensures the normal transmission of neuronal signals, thus effectively inhibiting nerve damage after hypoxia. In addition, there may be other mechanisms by which FA improves the learning and memory ability of rats with hypoxic-ischemic injury. Further research is needed to clarify the mechanism underlying the effect of FA in the treatment of HIBD and make better use of it in the clinic; traditional Chinese medicine containing FA provides a theoretical basis for the treatment of HIBD.

## Acknowledgments

We thank the Southwest Medical University Laboratory of Biochemistry and Molecular Biology for technical assistance and the Public Health Center for assistance and use of the Morris water maze. This work was supported by the Science and Technology Bureau of LuZhou City (2017LZXNYD-J30).

## Conflict of interest

The authors declare that there are no conflicts of interest.

